# NF-κB driven inflammation mediates loss of upper airway epithelial tolerance to *Streptococcus pneumoniae* during influenza co-infection

**DOI:** 10.1101/2025.10.25.684585

**Authors:** Zahrasadat Navaeiseddighi, Zhihan Wang, Kai Guo, Syed Hasan, Taylor Schmit, Jamanah Ahsan, Ramkumar Mathur, Junguk Hur, Nadeem Khan

## Abstract

*Streptococcus pneumoniae* asymptomatically colonizes the human nasopharynx, where epithelial tolerance maintains mucosal homeostasis. However, influenza A virus (IAV) co-infection transforms this tolerogenic state into an inflammatory environment that promotes bacterial outgrowth and invasion. Here, we identify a TGF-β1 dependent epithelial program that sustains mucosal tolerance during *Spn* colonization and demonstrate that IAV co-infection disrupts this pathway through IL-17RA-NF-κB driven inflammation in the nasopharynx. In a murine colonization model, TGF-β1 blockade enhanced pro-inflammatory cytokine production and neutrophil recruitment, resulting in inflammation-driven *Spn* clearance. IAV co-infection suppressed epithelial TGF-β1 signaling, increased TRAF6/NF-κB activation, and impaired tight junction integrity, leading to *Spn* dissemination. Mechanistically, IL-17RA signaling contributed to the hyperactivation of the TRAF6/NF-κB axis. Pharmacologic inhibition of TRAF6 or NF-κB restored epithelial barrier function and reduced *Spn* translocation in a human air-liquid interface nasopharyngeal epithelial model. These findings reveal a conserved epithelial signaling axis through which influenza disrupts mucosal tolerance and promotes *Spn* invasion, highlighting the canonical TRAF6-NF-κB pathway as a potential therapeutic target to preserve epithelial integrity and mitigate *Spn* infection during viral-bacterial co-infection of the upper respiratory tract.

## Introduction

*Streptococcus pneumoniae* (*Spn*) is a leading cause of non-invasive infections such as otitis media and pneumonia, and invasive pneumococcal disease (IPD), including bacteremia and meningitis [1,2]. *Spn* establishes asymptomatic colonization in the human nasopharynx (NP) across all age groups, with the highest and most persistent carriage observed in children (40– 60%) [3,4,5]. Importantly, *Spn* colonization induces low-grade, tolerogenic inflammation that promotes NP mucosal homeostasis while permitting gradual bacterial clearance [6,7]. Additionally, colonization serves as a natural immunizing event, promoting the development of antigen-specific mucosal and systemic immunity, including IgA production and memory T cell responses [8,9]. Paradoxically, while colonization induces protective immunity against subsequent *Spn* colonization events, it also predisposes hosts to *Spn* infection, particularly during respiratory viral co-infections such as influenza [10]. Experimental murine and human studies demonstrate that prior *Spn* colonization may predispose individuals to *Spn* infections upon viral co-infection, suggesting a shift from a tolerogenic to a pro-inflammatory mucosal environment, likely driven by altered epithelial-immune signaling [11,12,13]. However, the tolerogenic host factors that govern *Spn* persistence in the NP, and their disruption during viral co-infection, remain incompletely defined and represent an area of high clinical significance.

To define the host factors that promote mucosal tolerance to *Spn* colonization in NP, we profiled the nasal immune response in a murine model of stable *Spn* colonization. Among several immunoregulatory mediators screened, TGF-β1 emerged as the dominant anti-inflammatory cytokine associated with mucosal tolerance, whereas other mediators, including IL-33, IL-22, and IL-10, did not show similar associations. Antibody blockade of TGF-β1 accelerated bacterial clearance, establishing a functional role for TGF-β1 as a key regulator of mucosal tolerance to *Spn*. We next examined whether influenza co-infection alters the TGF-β1 driven tolerance program in NP. In *Spn* colonized mice, IAV co-infection suppressed TGF-β1 and disrupted TGF-β-SMAD3 signaling in NP epithelial cells. This was accompanied by a marked increase in TRAF6 and NF-κB activation, disruption of NP epithelial barrier integrity (reduced ZO-1), and enhanced *Spn* dissemination and invasion. This inflammatory shift was driven by IL-17RA dependent activation of TRAF6-NF-κB signaling in NP epithelial cells and was recapitulated in a human nasal ALI epithelial model, where pharmacologic inhibition of these pathways restored barrier integrity and reduced *Spn* translocation. Together, these findings reveal how viral co-infection disrupts the balance of epithelial tolerance and inflammation in the upper respiratory tract through dysregulation of the TGF-β1 and NF-κB signaling pathways, promoting *Spn* infection.

## Results

### TGF-β1 promotes mucosal tolerance to *Spn* colonization in the nasopharynx

The nasopharyngeal environment is evolutionarily adapted to maintain an anti-inflammatory state that permits bacterial colonization. However, the host mechanisms that sustain mucosal tolerance in the upper airway remain incompletely defined. To address this gap, we examined the expression of key anti-inflammatory mediators in nasopharyngeal mucosa from mice stably colonized with *Spn* serotype 6A. We focused on day 7 post-colonization, a time point associated with peak bacterial density, which provides an optimal window to assess steady-state host-microbe interactions. Western blot analysis revealed that, among several anti-inflammatory cytokines examined (IL-10, TGF-β1, IL-22, IL-33), TGF-β1 was the most strongly upregulated (Fig. 1A). Immunofluorescence staining of the nasal septa further confirmed robust TGF-β1 expression in response to *Spn* colonization (Fig. 1B). Since epithelial cells in NP are the primary site of bacterial colonization and key regulators of mucosal homeostasis, we hypothesized that epithelial TGF-β1 signaling underlies the establishment of the *Sp*n commensal state. To assess that, we performed western blot (pSMAD3) and immunofluorescence staining on NP epithelial cells (CD45^-^ Epcam^+^) cultured on chamber slides, showing robust pSMAD3 expression consistent with activation of the canonical TGF-β1 signaling pathway (Fig. 1C, D). To determine whether TGF-β1 signaling is essential for maintaining stable *Spn* colonization, we performed in vivo blockade using an anti-TGF-β1 antibody administered intraperitoneally starting on day 1 post-colonization and continued every other day until day 5 (for analysis at 7 dpi) or day 13 (for analysis at 14 dpi) (Fig. 1E). TGF-β1 blockade led to rapid *Spn* clearance from the nasopharynx, with a significantly reduced burden by day 7 and near-complete clearance by day 14, as determined by *Spn* CFUs in retrogradely collected nasal lavage. In contrast, control mice receiving isotype antibodies maintained stable colonization (Fig. 1F). Enhanced *Spn* clearance in TGF-β1 neutralized mice was accompanied by higher IL-1α and IL-6 protein levels (Fig. 1G) and increased neutrophil infiltration in NP lavage (Fig. 1H). Together, these findings identify TGF-β1 as a key regulator of epithelial tolerance that supports stable *Spn* colonization in the nasopharynx.

**Fig. 1.**
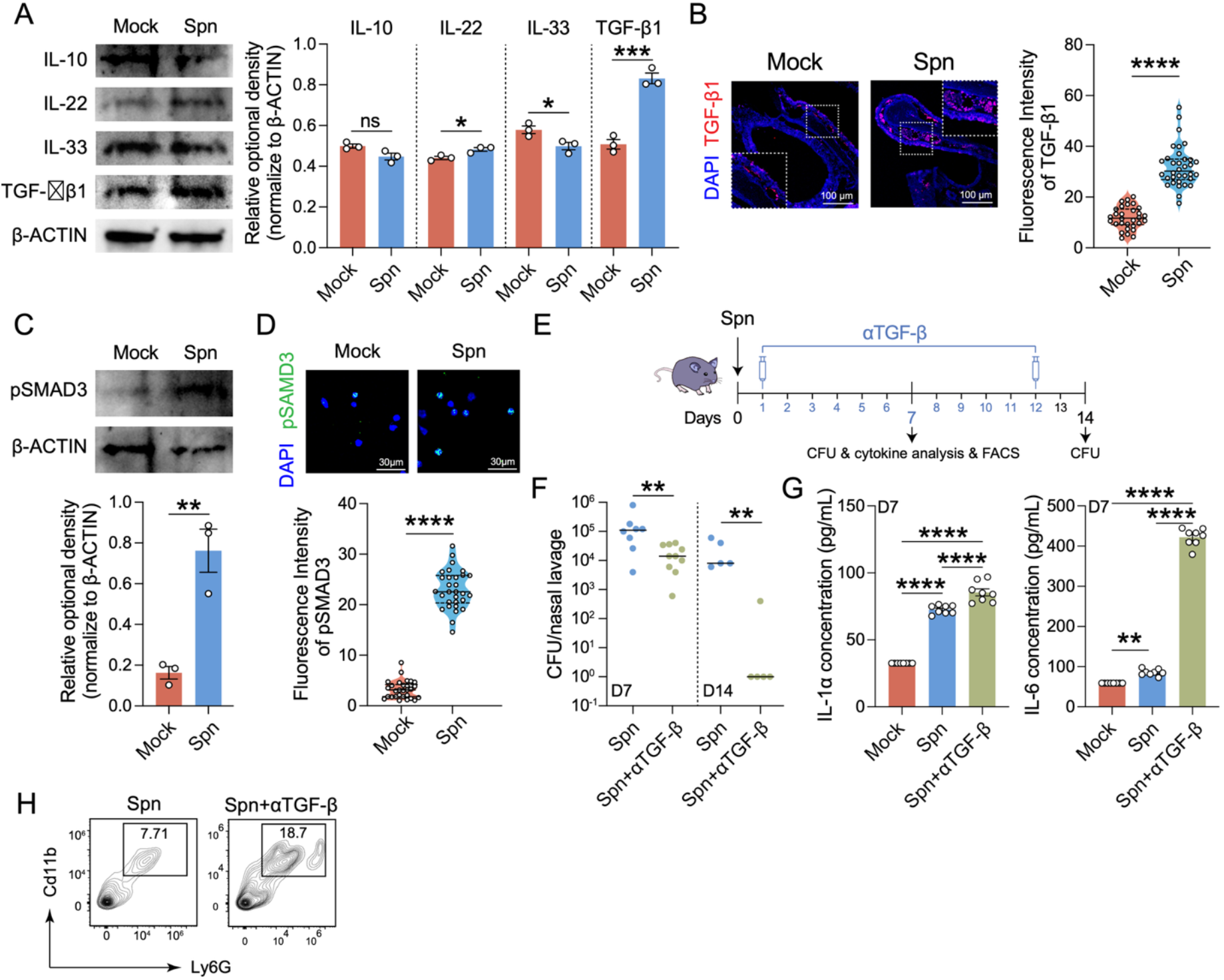
TGF-β1 regulates mucosal immune tolerance and promotes *Spn* colonization in NP. (A) Western blot analysis of tolerance-associated mediators (IL-10, IL-22, IL-33, TGF-β1) in nasopharyngeal tissues from mock and Spn-colonized mice at 7 dpi. (B) IF staining of nasal septa showing higher TGF-β1 expression within NP mucosa of Spn-colonized mice at 7 dpi. (C-D) Western blot and IF detection of pSMAD3 in sorted epithelial cells (CD45^−^ EpCAM^+^) from NP mucosa at 7 dpi. (E) Experimental design for in vivo TGF-β1 neutralization following Spn colonization (F). Mice were euthanized at 7 and 14-days post-infection to measure bacterial burden in the nasal lavage. (G) NP lavage from anti-TGF-β1 treated mice exhibited higher IL-6 and IL-1α levels, determined by flow cytometric bead array (H) Flow-cytometric quantification of neutrophils (Ly6G^+^ CD11b^+^) in NP lavage fluid upon TGF-β1 blockade. Data representative of at least two independent experiments (n=5 mice/group) were analyzed by One-Way ANOVA with Tukey’s post hoc. p < 0.05 (*), p < 0.01 (**), p < 0.001 (***), and p < 0.0001 (****) ns, not significant.

### IAV co-infection suppresses TGF-β1 mediated mucosal tolerance and amplifies TRAF6-NF-κB driven epithelial inflammation in the nasopharynx

Since TGF-β1 acted as a key mediator of mucosal tolerance in our *Spn* colonization model, we next investigated whether IAV co-infection disrupts this tolerance by suppressing TGF-β1 signaling in nasal epithelial cells. To test this, *Spn* colonized mice were co-infected with IAV 24 hours after colonization. Six days later, nasopharyngeal mucosal tissues were harvested, and epithelial cells were magnetically sorted as CD45^−^EpCAM^+^ populations. TGF-β1 protein levels in whole mucosal lysates were determined by western blot, while pSMAD3 expression was assessed by both western blot and immunofluorescence staining of sorted NP epithelial cells cultured on chamber slides. IAV co-infection reduced TGF-β1 protein levels in the nasal mucosa and SMAD3 phosphorylation in epithelial cells (Fig. 2A, B), indicating broad suppression of homeostatic TGF-β1 signaling in the nasopharynx. Prior studies [14], including our own [15,16], have shown that IL-17RA, IL-1, and TNF-α signaling drives mucosal epithelial inflammation via shared mediators such as TRAF6 and NF-κB. We therefore examined whether these signaling mediators are activated during co-infection and contribute to pathogenic epithelial inflammation.

**Fig. 2.**
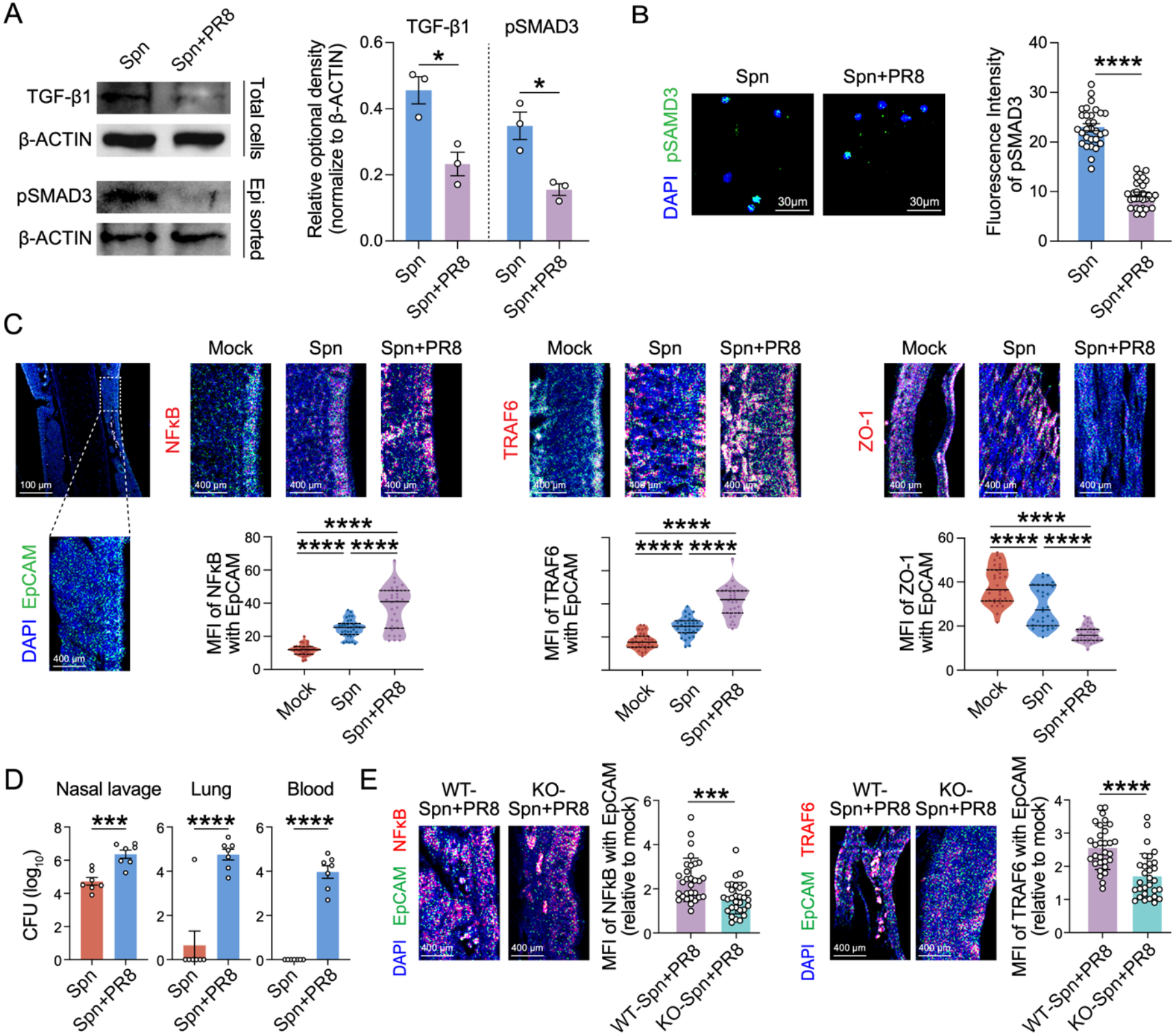
IAV co-infection disrupts TGF-β1-mediated epithelial tolerance and enhances NF-κB and TRAF6 activation in NP epithelial cells. (A) Western blot analysis showing reduced TGF-β1 (total homogenized nasopharyngeal mucosal tissues) and pSMAD3 (sorted epithelial cells: CD45-Epcam+ from NP tissues) during Spn+IAV co-infection compared to Spn alone at 6 days post co-infection. (B) IF staining of CD45-Epcam+ cells from nasopharyngeal mucosa showing decreased pSMAD3 expression during co-infection. (C) IF staining of nasal septa showing enhanced NF-κB (p65) and TRAF6 expression in NP epithelial cells (EpCAM^+^) following co-infection and reduced ZO-1 expression in NP epithelium of co-infected mice. (D) Spn CFUs in nasal lavage, lung, and blood showing increased Spn dissemination during co-infection compared to Spn colonization alone at 6 days post co-infection. (E) IF staining of nasal septa showing reduced NF-κB (p65) and TRAF6 expression in NP epithelial cells (EpCAM^+^) following KO-Spn+PR8 co-infection. Data are representative of at least two independent experiments (n= 5 mice per group). IF signals were quantified within predefined (unbiased) regions of interest (ROIs). Data were analyzed using one-way ANOVA with post hoc multiple-comparison test. Statistical significance is indicated as follows: p < 0.05 (*), p < 0.01 (**), p < 0.001 (***), and p < 0.0001 (****).

To address this, we assessed TRAF6 and NF-κB expressions in nasal septal epithelial cells by immunofluorescence staining, co-staining EpCAM with NF-κB (p65) or TRAF6. *Spn* colonization alone caused mild upregulation of TRAF6 and NF-κB, along with reduced ZO-1 expression, and these effects were markedly amplified during IAV co-infection (Fig. 2C). We further probed sorted epithelial cells for phosphorylated p65 and observed significantly higher phospho-p65 levels in co-infected epithelial cells, confirming enhanced NF-κB activation (Fig S1A). The increased epithelial NF-kB signaling correlated with higher *Spn* respiratory burden (NP lavage, lung) and invasion (blood) of co-infected mice (Fig. 3D). Next, we analyzed TRAF6 and NF-κB expression in nasal septal epithelial cells from WT and IL-17RA^−^/^−^ mice co-infected with *Spn* and PR8. IL-17RA-deficient mice showed a marked reduction in TRAF6 and NF-κB expression compared with WT mice (Fig. 2E). These findings are consistent with our previous report showing that IL-17RA deficiency reduces *Spn* pathogenesis and invasion in the nasopharynx, supporting a central role for IL-17RA dependent pathogenic signaling in driving epithelial inflammation and bacterial dissemination during IAV co-infection.

**Fig. 3.**
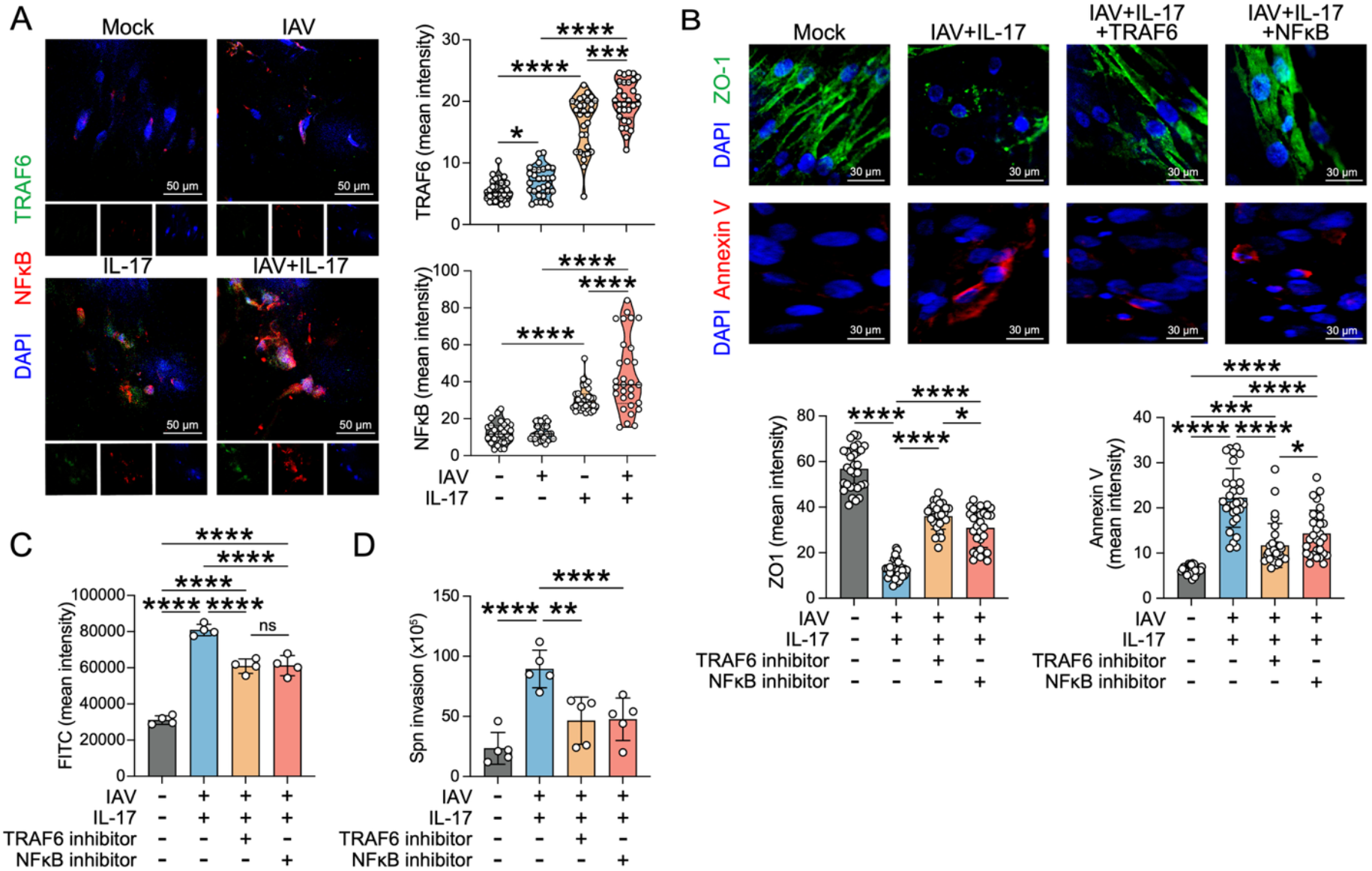
Therapeutic inhibition of TRAF6 and NF-κB signaling restores barrier integrity in a human ALI NP epithelial model. Primary human NP epithelial cells were cultured under ALI conditions for 21 days to allow differentiation, then infected with IAV (A/California/07/2009; MOI 0.1) and treated with recombinant IL-17A (200 ng/mL) in the presence or absence of TRAF6 (30uM) or NF-κB (10uM) inhibitors for 24 h. (A) IF staining showed increased TRAF6 and NF-κB expression in IL-17A-treated cells. B. IF staining shows increased ZO-1 expression and reduced Annexin V^+^ staining in inhibitor-treated cultures. (C) Quantification of FITC–dextran permeability across ALI cultures showing improved barrier upon TRAF6 and NF-κB inhibition. (D) Basolateral Spn translocation showing reduced bacterial passage across inhibitor-treated epithelium. Data are representative of at least two independent experiments (n = 3-4 independent ALI cultures per condition); IF signals were quantified within predefined regions of interest (ROIs). Data were analyzed using one-way ANOVA with post hoc multiple-comparison testing. Statistical significance is indicated as follows: p < 0.05 (*), p < 0.01 (**), p < 0.001 (***), and p < 0.0001 (****) ns, not significant.

### Pharmacologic blockade of TRAF6 or NF-κB signaling improves barrier integrity and attenuates *Spn* translocation in a human nasopharyngeal ALI epithelial model

Given that IL-17A associated TRAF6/NF-κB signaling correlated with epithelial inflammation and barrier disruption in vivo, we next used a human nasopharyngeal epithelial ALI model to test the functional impact of this pathway and determine whether its pharmacologic inhibition could restore epithelial integrity and reduce *Spn* translocation during IAV co-infection. To test this, primary human nasopharyngeal epithelial cells were grown under air-liquid interface culture as we previously reported [15]. Briefly, 30,000 cells were seeded onto transwell inserts and maintained under ALI conditions for three weeks to allow differentiation into a pseudostratified, ciliated epithelium. Differentiated ALI cultures were infected with a human IAV strain, A/California/07/2009 (H1N1) (MOI: 1) for 1 hour, followed by treatment with recombinant human IL-17A (200 ng/mL) and incubation for an additional 24 hours in the presence or absence of the TRAF6 inhibitor (10 μM: 18]) and the NF-κB inhibitor [5 μM; 17]. The impact of pathway inhibition on epithelial barrier integrity was evaluated by assessing epithelial apoptosis using Annexin V staining, ZO-1 expression, FITC-dextran permeability, and *Spn* translocation from the apical to the basolateral chamber. IL-17A treatment led to robust induction of TRAF6 and phosphorylated p65 expression in differentiated NP epithelial cultures (Fig. 3A), suggesting activation of the TRAF6-NF-κB pathway prior to inhibitor treatment. Inhibition of either TRAF6 or NF-κB individually led to a significant increase in ZO-1 expression and reduced Annexin V (Fig. 3B), which correlated with reduced basolateral transfer of FITC-dextran (Fig. 3C), indicating improved epithelial barrier integrity. To further assess barrier function, IAV-infected ALI cultures treated with cytokines and inhibitors were exposed to *Spn* (MOI: 1) for 1 hour, and basolateral translocation was quantified by plating basal chamber media on blood agar plates 2 hours post-infection. We observed markedly reduced *Spn* translocation to the basal chamber in inhibitor-treated cultures (Fig. 3D), consistent with higher ZO-1 expression, lower Annexin V expression, and decreased FITC-dextran leakage. Finally, TRAF6 inhibitor dramatically reduced IL-17A induced NF-κB (phospho-p65) expression (Fig. S1B), suggesting that TRAF6 functions as a key upstream mediator of NF-κB activation in nasopharyngeal epithelial cells. Collectively, these data provide functional evidence that IAV infection engages TRAF6 and NF-κB signaling to drive epithelial barrier disruption, and that targeted inhibition of this pathway ameliorates barrier integrity and limits *Spn* invasion.

## Discussion

Our findings define a previously unrecognized epithelial signaling axis that maintains mucosal tolerance during *Spn* colonization and demonstrate how IAV co-infection perturbs this balance through IL-17A-NF-κB mediated inflammation. The nasopharyngeal mucosa is constantly exposed to commensal and potentially pathogenic microbes, which requires a regulatory framework that limits tissue-damaging inflammation while maintaining barrier integrity. We show that TGF-β1 signaling is central to this equilibrium by limiting pro-inflammatory responses, thereby enabling a stable *Spn* colonization state. Antibody-mediated TGF-β1 blockade disrupted this homeostasis, resulting in increased pro-inflammatory cytokine production and neutrophil recruitment that promoted clearance of colonizing *Spn*. These findings identify TGF-β1 as a key determinant of epithelial tolerance that suppresses inflammation-driven bacterial clearance in the nasopharynx.

During IAV co-infection, this tolerogenic program was markedly suppressed. TGF-β1 and pSMAD3 were both reduced in nasopharyngeal tissue, correlating with higher TRAF6 and NF-κB activation and reduced epithelial ZO-1 expression, resulting in *Spn* dissemination into the lower respiratory tract and bloodstream. These observations support previous studies showing that viral infection promotes bacterial invasion by altering cytokine and epithelial TJ integrity [19,20,21]. Our findings demonstrate that viral-induced activation of the IL-17RA-NF-κB axis suppresses TGF-β1 signaling, thereby shifting the mucosal environment from a tolerogenic to an inflammatory state. IL-17RA deficiency suppressed TRAF6 and NF-κB activation in the nasopharyngeal epithelium, supporting a causal role for this pathway in IAV-driven inflammation in the upper airway. In our prior work with the *Spn*-IAV co-infection model, we showed that IL-17RA deficiency markedly reduced *Spn* pathogenesis and invasion in the nasopharynx [15], suggesting that IL-17RA dependent signaling promotes epithelial inflammation and bacterial dissemination. The current findings extend that observation by defining the downstream TRAF6-NF-κB pathway as a mechanistic link between IL-17RA signaling and epithelial barrier disruption during viral co-infection. Thus, the interplay between TGF-β1 dependent tolerance and IL-17RA driven inflammation defines the outcome of airway epithelial responses and *Spn* pathogenesis during *Spn*-IAV co-infection in the upper respiratory tract.

Supporting these in vivo findings, our data further show that TRAF6-NF-κB activation is a driver of epithelial barrier disruption in an ALI model of human NP epithelial cells. IAV infection and IL-17A treatment of differentiated human nasopharyngeal epithelial cultures reproduced the in vivo phenotype, inducing epithelial apoptosis, loss of ZO-1, and increased barrier permeability that facilitated *Spn* translocation. Pharmacologic inhibition of TRAF6 or NF-κB reduced epithelial apoptosis and restored tight junction integrity, leading to reduced *Spn* translocation. These findings are consistent with previous studies in airway and intestinal epithelia showing that while NF-κB signaling is essential for maintaining tight junction and barrier integrity under homeostatic conditions, its inhibition enhances tight-junction stability during inflammatory stress, highlighting a context-dependent dual role [22,23]. Therapeutically, these findings suggest that selective inhibition of TRAF6 or NF-κB signaling within the inflamed epithelium could represent a targeted strategy to preserve barrier integrity and limit *Spn* invasion without broadly suppressing host immunity. These data support a model in which IAV co-infection drives IL-17RA dependent NF-κB hyper-activation, which suppresses TGF-β1 mediated tolerance to promote epithelial inflammation and *Spn* dissemination in the nasopharynx. Collectively, these findings underscore the central role of epithelial signaling in maintaining barrier integrity and reveal how its dysregulation by viral co-infection can set the stage for secondary bacterial invasion, providing avenues to target epithelial-intrinsic inflammation to curb severe *Spn* infection during IAV.

## Materials and methods

### Ethics statement

C57BL/6 mice were purchased from Jackson Laboratory and bred in-house at the University of Florida in accordance with the UF Institutional Animal Care and Use Committee guidelines. IL-17RA^−^/^−^ mice were provided by Sarah Geffen (University of Pittsburgh, PA) and are proprietary to Amgen (Thousand Oaks, CA). All animal procedures were conducted under the University of Florida Institutional Animal Care and Use Committee protocol #IACUC202100000057.

### Animals and Microbial Strains

An equal number of age-matched (6-8-week old) male and female mice were included in the study. The mouse-adapted influenza A H1N1 A/Puerto Rico/8/1934 (IAV or PR8) virus [24] was purchased from Charles River. The human IAV H1N1 pdm09 strain A/California/04/2009 was obtained from the American Type Culture Collection (ATCC). *Spn* serotype 6A strain BG7322 was provided by Rochester General Hospital Research Institute and has been extensively used by our group [25,15,26].

### Infection models

Mice were anesthetized with isoflurane (4% v/v) and inoculated (i.n.) with *Spn* 6A (BG7322, 1×10^5^ CFU). Colonized mice were infected with PR8 at 1000 PFU, 24 hours later. Both *Spn* and PR8 inocula were administered in 10 μL PBS intranasally. Control mice received 10 μL PBS. Mice were euthanized at 7 and 14 dpi (colonization) or 6 days post co-infection. Retrograde NP lavage was collected using 200 μL PBS. Lung homogenates were aseptically isolated, and blood was collected via cardiac puncture. Serially diluted NP lavage, blood, and homogenized lungs were plated on blood agar, and CFUs were counted the next day.

### Flow cytometry

NP lavage samples were first pooled, and 100 μL of diluted Fc block was added to each lavage and incubated for 20 min on ice. The samples were subsequently stained with antibodies against CD11b (PE-Cy7) and Ly6G (APC) for 30 min at room temperature in the dark. A Cytek Aurora Flow Cytometer was used to acquire 30,000 events per lavage sample. Data were analyzed using FlowJo v10. Cytokine quantification in NP lavage was performed using the LEGENDplex™ Mouse Inflammation Panel Kit (13-plex) (Cat. No. 740446). Samples were acquired on an Attune™ NxT Acoustic Flow Cytometer and analyzed using LEGENDplex™ Data Analysis Software (BioLegend).

### IF staining and quantification

Cells were prepared from the NP mucosal tissue of Mock, *Spn* (7 days post infection), and *Spn*+PR8 co-infection group (6 days post co-infection). Epithelial cells were magnetically sorted using CD45 (CD45 MicroBeads, Cat# 130-052-301, Miltenyibiotec) and Epcam (CD326: EpCAM Microbeads, Cat#130-105-958, Miltenyibiotec). Sorted epithelial cells were centrifuged at 250xg, resuspended in PBS (1×10^6^/ml), and adhered to coverslips for 30 mins at RT. Adhered cells were fixed (4% paraformaldehyde), permeabilized (0.1% Triton X-100), blocked (10% donkey serum), stained with anti-pSMAD3 (Abcam: ab52903) overnight, followed by secondary antibody and nuclear staining with DAPI. Paraffin-embedded nasopharyngeal sections from mock, *Spn, Spn*+PR8, and IL-17RA KO-*Spn*+PR8 mice were stained with primary antibody EpCAM (Catalog# 13-5791-82, Thermo-fisher), TGFβ1 (Catalog# MA5-18023, Thermo-fisher) NFkB/p65 (Cat# ab16502, Abcam), phospho p65 (.Cell Signaling: #3033)), TRAF6 (Cat # sc8409, Santa Cruz), and ZO-1 (Cat# ab221547), followed by incubation with secondary antibody and counter-staining with DAPI. The images were captured using a Zeiss 710 confocal laser scanning microscope at the Center for Immunology and Transplantation, University of Florida. The image analysis was done with NIH Fiji (ImageJ) software, and fluorescence intensity was quantified within defined regions of interest (ROIs) for each image.

### Western blot

NP mucosal tissues from mock, *Spn* colonized, and *Spn*+PR8 co-infected mice were collected at 7 dpi, and NP epithelial cells were magnetically sorted using CD45 and Epcam beads (CD45^-^ Epcam^+^). Tissues were homogenized, and magnetically sorted epithelial (CD45^-^ Epcam^+^) cells were lysed with RIPA lysis buffer with a protease and phosphatase inhibitor mixture (ThermoFisherScientific). Samples were centrifuged at 14,000xg for 10 min at 4°C, and supernatants were aliquoted and preserved at -80°C. Twenty micrograms of protein lysates were used for the Western blot and probed with TGF-β1 (Catalog# MA5-18023, Thermo-fisher), phospho p65 (ab76302), or pSMAD3 (Abcam: ab52903).

### Expansion and differentiation of HNEpCs under ALI culture and Immunostaining assay

Primary human nasal epithelial cells (HNEpCs, PromoCell: C-12620) were seeded on Transwell (Corning, Cat# 3470) at 35,000 cells per well and cultured under submerged conditions in PromoCell growth media. After 3-4 days, cells were differentiated with PneumaCult-ALI media (Cat#05001, StemCell) for 21 days, with basal media changed every 2 days. Cells were infected with the H1N1 pdm09 strain (MOI: 1) for 2 hours, washed with DPBS, and stimulated with recombinant human IL-17A (200 ng/mL) in the presence or absence of IAV in ALI medium for 24 hours. Finally, cells were fixed and stained with NFκB and TRAF6, followed by secondary antibodies and DAPI staining for confocal microscopy.

### Therapeutic inhibition of NFkB, TRAF6, and Analysis of epithelial barrier Integrity

At day 21 of ALI culture, the monolayer of differentiated epithelial cells was washed with DPBS (stem cell technologies). Cells were treated with NFkB inhibitor at 30uM (Cat#JSH-23, MCE MedChemExpress) and TRAF6 inhibitor at 10uM (Catalog No.: A18705, Adooq bioscience), followed by fixation and staining with tight junction protein ZO-1 (Cat# ab 221547, Abcam), Annexin-V (Cat# 66245-1-IG, Protein tech) as described for the immunofluorescence microscopy. FITC-dextran (3 mg/mL, 200 μL) was added to the apical chamber of the Transwell, and ALI maintenance media was added to the basal chamber and incubated for 2 hours at 37°C. Basal media (50 μL) was collected, diluted serially, and fluorescence measured (Ex: 490 nm/Em: 520 nm). For the *Spn* membrane translocation assay, *Spn* 6A (BG7322, MOI: 1) was added to the apical chamber in antibiotic-free DMEM for 2 hours at 37^0^C in a CO2 incubator [27]. Media was collected from the basal chamber, serially diluted, plated on blood agar, and *Spn* bacterial colonies counted on the next day.

## Acknowledgements

This work was supported by NIH grants R01 AI143741 and R21 AI151522 to N.K. The UND Flow Cytometry Core Facility was supported by NIH COBRE grant 5P20GM113123 and INBRE grant 5P20GM103442.

---

**Figure S1.**
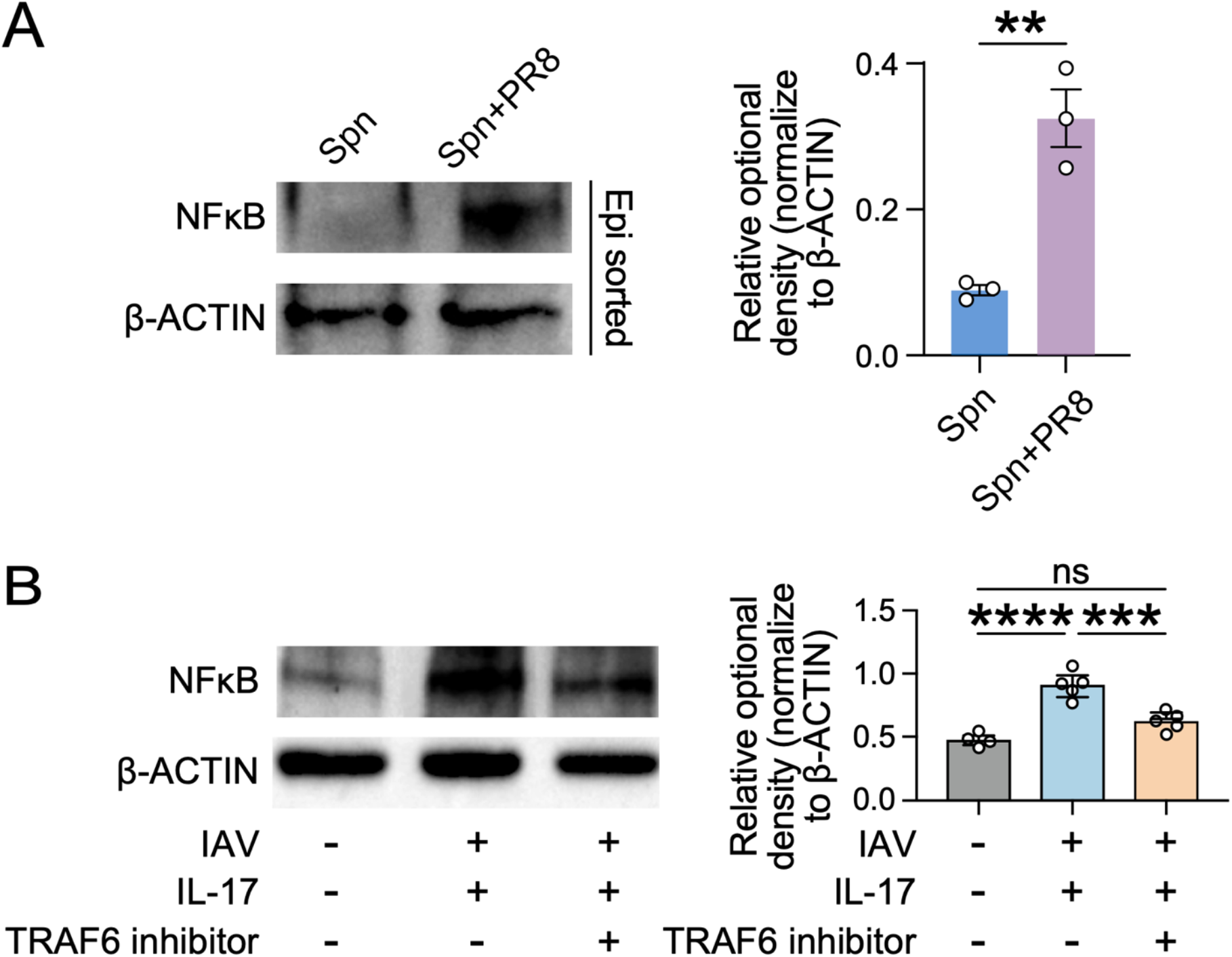
NF-κB activation and TRAF6 dependency in nasopharyngeal epithelial responses. (A) Western blot analysis showing increased NF-κB activation (phospho p65) in magnetically sorted epithelial cells from nasopharyngeal mucosa of *Spn* colonized and *Spn*+PR8 co-infected mice. Densitometric quantification (right) shows relative phospho p65 protein levels normalized to β-ACTIN. Data represent mean ± SEM. p < 0.01 (**). (B) Immunoblot showing NF-κB expression in human nasopharyngeal ALI cultures treated with IL-17A (200 ng/mL) and/or IAV (MOI 0.1) with or without TRAF6 inhibitor (10 µM). TRAF6 inhibition markedly reduced NF-κB activation (phospho p65) induced by IL-17A+IAV stimulation. Quantification (right) shows relative NF-κB expression normalized to β-ACTIN (mean ± SEM; *p* < 0.05; ns, not significant). p < 0.0001 (****). ns, not significant.

